# A new NHGRI Sample Repository for Human Genetic Research collection of induced pluripotent stem cell lines

**DOI:** 10.1101/2025.08.05.668740

**Authors:** Tatyana Pozner, Christine Grandizio, Matthew W Mitchell, Xiaoxia Cui, Amber Neilson, Monica F. Sentmanat, Ting Wang, Nathan O Stitziel, Nahid Turan, Laura B Scheinfeldt

## Abstract

We describe here a new NHGRI Sample Repository for Human Genetic Research collection of induced pluripotent stem cell (iPSC) lines reprogrammed from whole blood derived peripheral blood mononuclear cells (PBMCs). PBMCs were reprogrammed using Sendai viral vectors carrying transcription factors OCT4, SOX2, KLF4, and c-MYC. All iPSC lines exhibit a normal karyotype, express common stemness and pluripotency markers, and demonstrate the ability to differentiate into cell types representing all three germ layers. This iPSC collection (n=7) will have accompanying public, near telomere to telomere genomic data through the Human Pangenome Reference Consortium, and provides an invaluable new *in vitro* resource for studying common genetic and genomic variation and its functional implications.

## Introduction

The NHGRI Sample Repository for Human Genetic Research (NHGRI Repository) housed at the Coriell Institute for Medical Research (Coriell) facilitates studies of human genetic and genomic variation by establishing, characterizing and distributing a large (n>3,700) publicly available collection of renewable biospecimens donated by communities living around the world, including the communities that have participated in the 1000 Genomes Project and the Human Pangenome Reference Consortium ^1-3^. NHGRI Repository participants have consented to their biospecimens being used for a wide range of general research and to public data sharing of large-scale genomic data collected from their samples.

This new collection of seven induced pluripotent stem cell (iPSC) lines was generated from peripheral blood samples collected as part of a larger effort to improve the human reference genome to better represent human genetic variation ^2,4,5^. This collection is an important new resource to improve our understanding of tissue-specific gene regulation and cellular function.

## Results

We reprogrammed seven iPSC lines (NHGRI Repository identifiers: HG06799, HG06800, HG06801, HG06802, HG06803, and HG06804, HG06807) from peripheral blood mononuclear cells (PBMCs) using the CytoTune 2.0 Sendai Reprogramming Kit (Thermo Fisher Scientific), which includes the pluripotency factors OCT4, SOX2, KLF4, and c-MYC. Five of the seven iPSC lines in the collection were reprogrammed at Coriell (HG06800, HG06801, HG06802, HG06803, and HG06804), and two of the seven iPSC lines in the collection were reprogrammed at Washington University (HG06799, HG06807). All iPSCs in the collection passed all quality control assessments ^6^ (**Table 1**): they display typical iPSC morphology under phase contrast microscopy, exhibit alkaline phosphatase activity (**Figure S1**), and are positive for Stage-Specific Embryonic Antigen 4 (SSEA-4; **Figure 1A**). In addition, the expression of several pluripotency markers was evaluated in cell lines reprogrammed at Coriell, using an immunocytochemical assay (**Figure 2**). We assessed post-thaw cell viability by culturing a frozen vial, with iPSC colony area increasing 6-28-fold over a 3-5 day period (**Table S1**). We performed cytogenomic G-banding analysis to confirm normal karyotypes (**Figure 1B**). We did not detect any Sendai virus (SeV) genome or transgenes by qRT-PCR using SeV-specific primers (**Table S2**) after passage 10 in the five iPSCs reprogrammed at Coriell. In addition, we did not detect any SeV genome or transgenes by qRT-PCR using SeV-specific primers (**Table S2**) in the two iPSCs reprogrammed at Washington University. We confirmed pluripotency by an embryoid body (EB) formation assay (**Figure 1C**). The lines are free of mycoplasma contamination **(Table 1, Table S3**), and microsatellite profiling has confirmed iPSC authenticity (**Table 1, Table S3**).

**Table 1.** Quality Control characterization and validation of iPSCs reprogrammed at Coriell.

| **Classification** | **Test** | **Result** |
| --- | --- | --- |
| **Morphology** | Photography | Normal |
| **Phenotype** | Quantitative analysis (i.e. Flow cytometry) | SSEA-4 (94.78% - 99.22%) |
| **Genotype** | Karyotype (G-banding and microarray) and resolution | HG06800 – 46, XX<br>HG06801 – 46, XY<br>HG06802 – 46, XX<br>HG06803 – 46, XX<br>HG06804 – 46, XY |
| **Identity** | MSAT analysis | Six sites tested, all sites matched |
| **Microbiology and virology** | Mycoplasma | Mycoplasma testing by qRT-PCR. Negative |
| **Differentiation potential** | Embryoid body formation | Embryoid body (EB) with three germ layers formation (ectoderm, mesoderm, and endoderm) |

**Figure 1.**
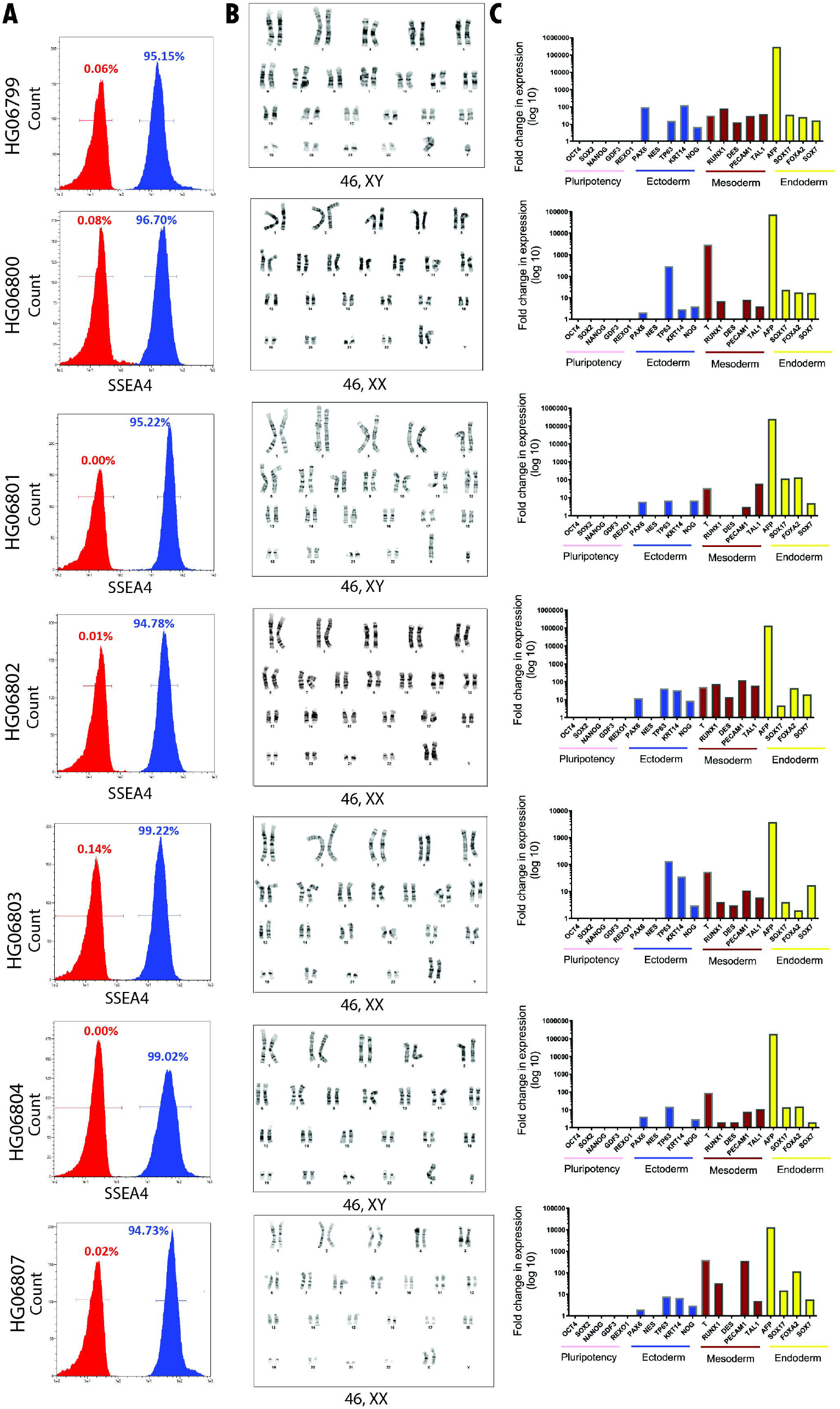
Characterization and quality control assessment of induced pluripotent stem cells (iPSCs). A) Stage specific embryonic antigen 4 (SSEA4) staining, B) G-banding karyograms. C) Fold change in expression of pluripotency genes and tri-lineage specific genes between undifferentiated cells and cells differentiated by embryoid body formation.

**Figure 2.**
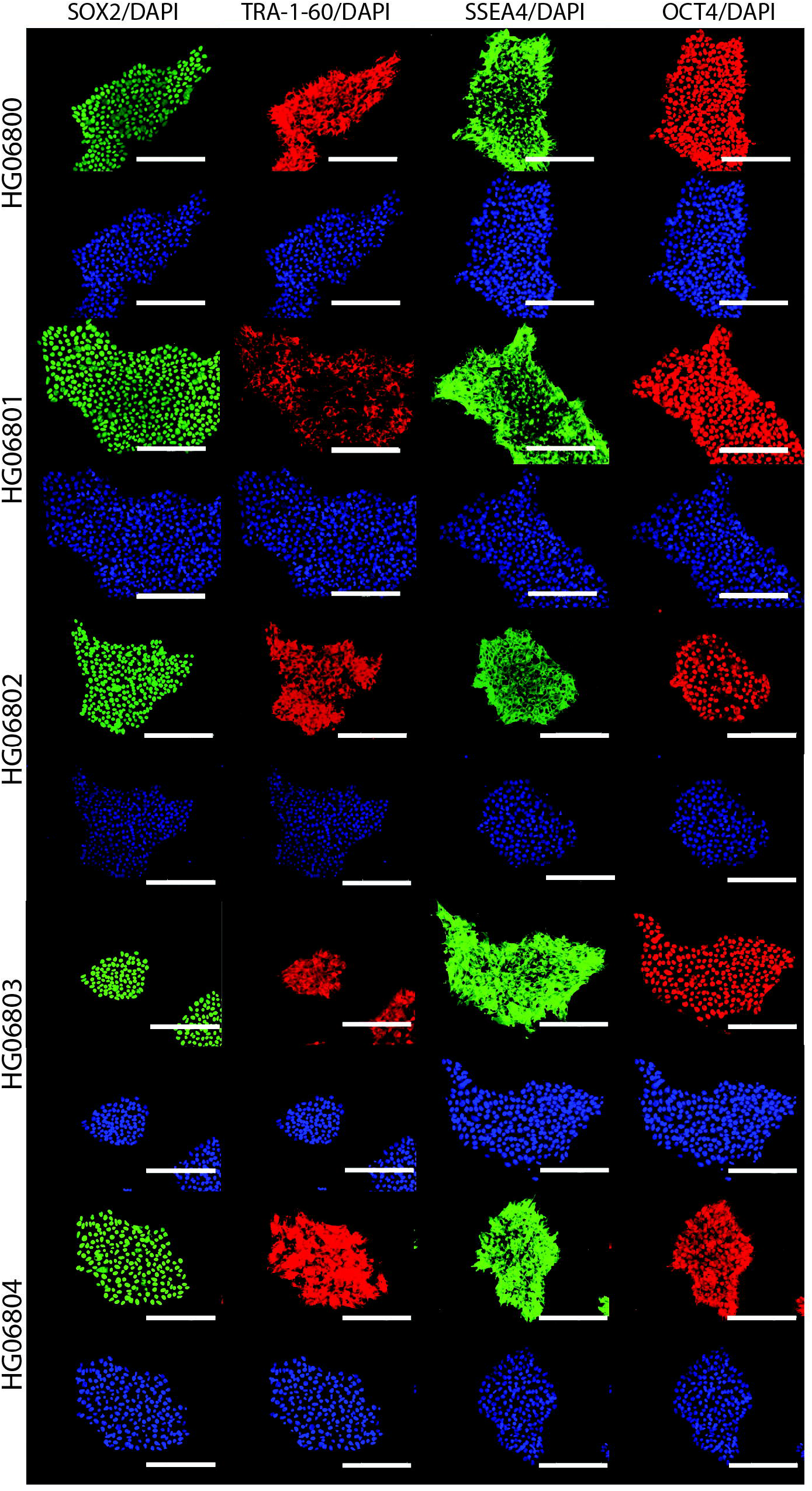
Pluripotency assessment of cell lines reprogrammed at Coriell. Example immunofluorescence images showing expression of key pluripotency markers SOX2, OCT4, SSEA4, and TRA-1-60 in iPSCs. Scale bar: 200 μm.

## Methods

### Samples

The samples included in the new NHGRI Repository iPSC resource were collected as part of a larger effort to improve the human reference genome with more comprehensive coverage of the genome, as well as to increase representation of human genetic variation ^2,4,5^. The participants that generously agreed to donate to the repository consented to the creation of immortalized cell lines, including iPSCs, to their cell lines and DNA being made available for general research usage, and to large-scale genomic data collected from their samples being made public. In addition, these samples are in the process of being characterized by the HPRC with near telomere to telomere genomic assemblies, and these public data will add important genetic and genomic characterizations to this iPSC collection. The recommended language for referring to these samples is: African Americans living in St. Louis, Missouri, which may be abbreviated to ASL after the full description is provided (https://catalog.coriell.org/1/NHGRI/About/Guidelines-for-Referring-to-Populations).

### Reprogramming and Cell Culture

The five iPSCs reprogrammed at Coriell were reprogrammed from PBMCs using CytoTune 2.0 (Thermo Fisher Scientific) Sendai vectors encoding OCT4, SOX2, KLF4, C-MYC, and EmGFP. One day after transduction, we cultured cells in complete StemPro™-34 medium (ThermoFisher Scientific) supplemented with L-Glutamine (2mM), CSF (100 ng/ml), FLT-3 (100 ng/ml), IL-3 (20 ng/ml) and IL-6 (20 ng/ml) and replated them on day 3 post transduction on mouse embryonic fibroblasts (MEF) coated culture dishes. From day 3 through day 7 post-transduction, the medium was replaced every other day with fresh complete StemPro™-34 medium without cytokines. On day 7 post-transduction, half of the culture medium was replaced with iPSC culture medium consisting of DMEM/F12 (Gibco) supplemented with 20% (v/v) KnockOut Serum Replacement (Invitrogen), 2 mM L-glutamine (Invitrogen), 1× non-essential amino acids (Invitrogen), 55 µM 2-mercaptoethanol (Invitrogen) and bFGF (10ng/mL). From day 8 onward, the culture medium was fully replaced daily. After 2-3 weeks post transduction, we selected 24 colonies and expanded them. At passage 10, we cryopreserved the lines. Cells were transitioned to Matrigel and maintained in mTeSR1 prior to incorporation into the NHGRI Repository.

The two iPSCs reprogrammed at the Genome Engineering & Stem Cell Center (GESC@MGI) at Washington University were reprogrammed from erythroid progenitors enriched and expanded from PBMCs in StemSpan SFEM II medium (STEMCELL Technologies) with a StemSpan Erythroid expansion supplement (STEMCELL Technologies). The progenitors were cultured in this medium on the day of transduction using CytoTune 2.0 (Thermo Fisher Scientific) and transferred to a 1:1 ratio of erythroid medium:mTeSR+ (STEMCELL Technologies). On day 3 post transduction, the cells were transferred to 1:3 ratio of erythroid medium:mTeSR+. On day 4 post transduction, cells were transferred to 100% mTeSR+ and plated on a Matrigel-coated surface. After 2-3 weeks,12 colonies were selected, and three clones with the best morphology were expanded at passage 5 and subsequently cryopreserved.

We thawed all cryopreserved cells by warming at 37°C, tested them for sterility, and cultured them on Matrigel-coated plates in mTeSR1 supplemented with ROCK inhibitor for 24 hours. Media were then changed daily. We passaged the cells at 75-85% confluency using Versene (2 minutes and room temperature) or ReLeSR (5-7 min, 37°C).

### Alkaline Phosphatase Staining

We stained iPSCs using the StemTAG™ Alkaline Phosphatase Staining Kit (Cell Biolabs, Inc.).

### Immunocytochemistry

We characterized the cells using a PSC Immunocytochemistry Kit with SOX2/TRA-1-60 and SSEA4/OCT4 antibody pairs. After fixation, permeabilization, and blocking, we applied primary antibodies (3h, 4°C), followed by secondary antibodies (1h, RT). We added DAPI during the final PBS wash (5 min).

### Flow Cytometry

We quantified surface antigen expression of iPSC markers by flow cytometry. We dissociated the cells with Trypsin, washed the cells with PBS, and incubated the cells with fluorophore-conjugated antibodies (Table S2) for 15 min at room temperature. We used the MACSQuant Flow Cytometer and MACSQuantify software for all related analyses.

### G-Banded Karyotyping

Twenty metaphase cells were counted and examined, and five selected metaphase cells were karyotyped.

### Cell Line Authentication

We assessed cell line identity and authenticity by extracting DNA from each line and comparing the microsatellite (MSAT) marker profile to the DNA extracted from the whole blood sample submission as previously described ^7^.

### Mycoplasma Detection and Sterility Testing

We tested for mycoplasma contamination using the MycoSEQ™ Mycoplasma Detection Kit, a real-time PCR assay detecting over 90 species with <10 copies sensitivity. We assessed cell culture sterility via growth assays on trypticase soy agar and Sabouraud dextrose.

### Sendai Virus Detection

We extracted total RNA using the RNeasy Plus Mini Kit. We synthesized cDNA from RNA using the High Capacity cDNA Reverse Transcription Kit. We quantified gene expression by qRT-PCR (primers used at Coriell and primers used at GESC@MGI are listed in **Table S2**).

### Differentiation Potential

We assessed pluripotency via embryoid body (EB) formation. We cultured EBs in DMEM with 10% FBS, 1% L-glutamine, NEAA, and sodium pyruvate for 10 days. We extracted RNA using the RNeasy Mini Kit and quantified RNA by NanoDrop One. We synthesized cDNA, analyzed gene expression by qRT-PCR, and normalized Ct values to GAPDH. We calculated relative expression as fold change compared to undifferentiated cells using the Livak-Schmittgen method ^8^.

## Discussion

Here, we introduce a new collection of seven iPSCs that have been incorporated into the NHGRI Repository. These cell lines were created from whole blood samples donated by African Americans living in St. Louis, MO. The iPSC lines are currently available to the research community through the NHGRI Repository, housed at the Coriell Institute for Medical Research (https://www.coriell.org/1/NHGRI).

A distinctive feature of this collection is that each iPSC line has a corresponding lymphoblastoid cell line (LCL), and DNAs extracted from these LCLs are in the process of being extensively genome sequenced through the Human Pangenome Reference Consortium; telomere-to-telomere (T2T) assemblies for a subset of three of these LCLs are publicly available through related efforts ^3^ (https://github.com/biomonika/HPP/tree/main/T2T-Pedigree-project%20). All associated genomic data will be publicly accessible and will include high-quality, robust, long-read, near telomere-to-telomere (T2T) assemblies ^2^. The availability of matched iPSC and LCL lines enables comparative studies of cellular phenotypes and lineage-specific gene regulation. Moreover, the accompanying genomic data will facilitate a more comprehensive investigation of genetic and genomic variation and its functional consequences.

This resource will be particularly valuable for investigating tissue-specific gene regulation, cellular differentiation pathways, and the functional consequences of genetic and genomic variation on cellular function. Furthermore, these well-characterized lines will serve as important reference materials for future studies involving iPSCs.

