## Supplemental Material for "A new NHGRI Sample Repository for Human Genetic Research collection of induced pluripotent stem cell lines"

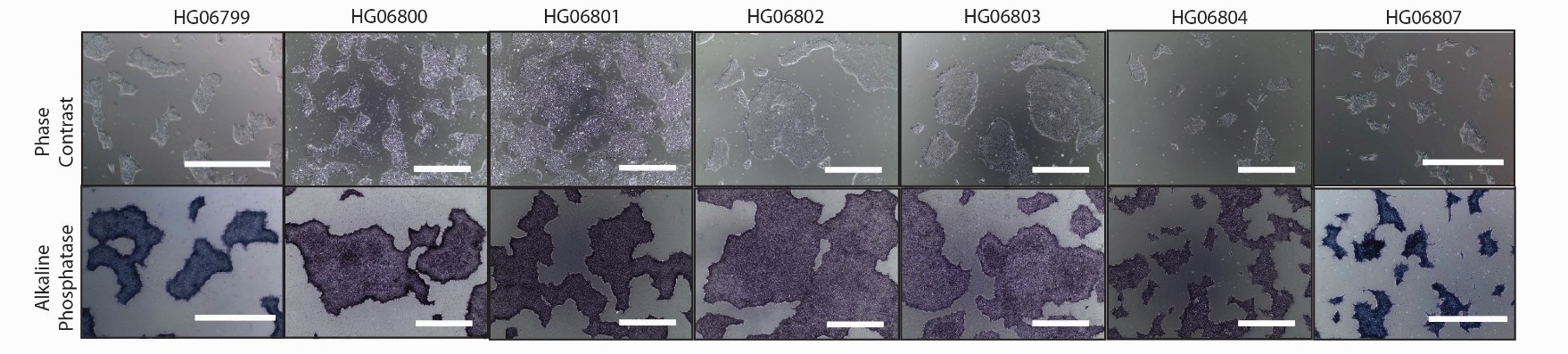
**Supplementary materials**

**Figure S1**. Quality control assessments of cellular morphology and pluripotency. The morphology of iPSCs was assessed under a phase contrast microscope. iPSC colonies exhibit alkaline phosphatase activity (purple), demonstrating maintained pluripotency. Scale bar = 1000 μm.

**Supplementary Table 1**. Post-thaw viability

| Cell line | Day | Average area (µm²) | Fold increase |
| --- | --- | --- | --- |
| HG06799 | 1 | 9,673 | 6 |
|  | 3 | 58,851 |  |
| HG06800 | 1 | 11,566 |  |
|  | 3 | 102,024 | 9 |
| HG06801 | 1 | 30,675 |  |
|  | 3 | 289,990 | 9 |
| HG06802 | 1 | 7,043 |  |
|  | 5 | 197,699 | 28 |
| HG06803 | 1 | 10,471 |  |
|  | 4 | 253,646 | 24 |
| HG06804 | 1 | 11,122 |  |
|  | 5 | 132,754 | 12 |
| HG06807 | 1 | 3,154 |  |
|  | 3 | 47,320 | 15 |

**Supplementary Table 2**. Reagents details

| **Antibodies used for immunocytochemistry/flow-cytometry** | | | |
| --- | --- | --- | --- |
|  | **Antibody** | **Company Cat #** | **RRID** |
| Pluripotency Markers | Alexa Fluor 647 anti human SSEA4 | BioLegend Cat# 330408 | RRID:AB_1089200 |
| Pluripotency Markers | Alexa Fluor 647 Mouse IgG3 κ | BioLegend Cat# 401321 | RRID: AB_10683445 |
| Pluripotency Markers | Isotype Control Alexa Fluor 488 anti-human/mouse SSEA3 | BioLegend Cat# 330306 | RRID: AB_1279440 |
| Pluripotency Markers | Alexa Fluor 488 Rat IgM κ Isotype Control | BioLegend Cat# 400811 | RRID:AB_1659271 |
| Primary Antibody | anti-SOX2 (host: rat) | ThermoFisher Scientific Cat# A24759 | RRID:AB_2651000 |
| Primary Antibody | anti-TRA-1–60 (host: mouse IgM | ThermoFisher Scientific Cat# A24868 | RRID:AB_2651002 |
| Primary Antibody | anti-OCT4 (host: rabbit) | ThermoFisher Scientific Cat# [A24867](https://www.ncbi.nlm.nih.gov/protein/A24867) | RRID:AB_2650999 |
| Primary Antibody | anti-SSEA4 (host: mouse IgG3) | ThermoFisher Scientific Cat# [A24866](https://www.ncbi.nlm.nih.gov/protein/A24866) | RRID:AB_2651001 |
| Secondary Antibody | Alexa Fluor 488 donkey anti-rat | Thermo Fisher Scientific Cat# A24876 | RRID:AB_2651007 |
| Secondary Antibody | Alexa Fluor 555 goat anti-mouse IgM | Thermo Fisher Scientific Cat# A24871 | RRID:AB_2651009 |
| Secondary Antibody | Alexa Fluor 555 donkey anti-rabbit | Thermo Fisher Scientific Cat# [A24869](https://www.ncbi.nlm.nih.gov/protein/A24869) | RRID:AB_2651006 |
| Secondary Antibody | Alexa Fluor 488 goat anti-mouse IgG3 | Thermo Fisher Scientific Cat# A24877 | RRID:AB_2651008 |
| **Primers** | | | |
|  | **Target** | **Forward/Reverse primer (5′-3′)** | |
| Sendai virus test (qPCR) – Coriell | SEV | Mr04269880_mr (TaqMan® probe ID) | |
| Sendai virus test (qPCR) - Coriell | SEV-KOS | Mr04421257_mr (TaqMan® probe ID) | |
| Sendai virus test (qPCR) - Coriell | SEV-KLF4 | Mr04421256_mr (TaqMan® probe ID) | |
| Sendai virus test (qPCR) - Coriell | SEV-CMYC | Mr04269876_mr (TaqMan® probe ID) | |
| Sendai virus test (qPCR) – Washington University | SEV | Forward: GGA TCA CTA GGT GAT ATC GAG C  Reverse: ACC AGA CAA GAG TTT AAG AGA TAT GTA TC | |
| Sendai virus test (qPCR) – Washington University | SEV-KOS | Forward: ATG CAC CGC TAC GAC GTG AGC GC  Reverse: ACC TTG ACA ATC CTG ATG TGG | |
| Sendai virus test (qPCR) – Washington University | SEV-KLF4 | Forward: TTC CTG CAT GCC AGA GGA GCC C  Reverse: AAT GTA TCG AAG GTG CTC AA | |
| Sendai virus test (qPCR) – Washington University | SEV-CMYC | Forward: TAA CTG ACT AGC AGG CTT GTC G  Reverse: TCC ACA TAC AGT CCT GGA TGA TGA TG | |
| Pluripotency Markers (qPCR) | OCT4 | hs00742896_s1 (TaqMan® probe ID) | |
| Pluripotency Markers (qPCR) | SOX2 | hs00602736_s1 (TaqMan® probe ID) | |
| Pluripotency Markers (qPCR) | NANOG | hs02387400_g1 (TaqMan® probe ID) | |
| Pluripotency Markers (qPCR) | GDF3 | hs00220998_m1 (TaqMan® probe ID) | |
| Pluripotency Markers (qPCR) | REXO1 | hs00381890_m1 (TaqMan® probe ID) | |
| House-Keeping Gene (qPCR) | GAPDH | hs00266705_g1 (TaqMan® probe ID) | |
| Differentiation Markers (qPCR) | PAX6 | hs00240871_m1 (TaqMan® probe ID) | |
| Differentiation Markers (qPCR) | NESTIN | hs00707120_s1 (TaqMan® probe ID) | |
| Differentiation Markers (qPCR) | TP63 | hs00978340_m1 (TaqMan® probe ID) | |
| Differentiation Markers (qPCR) | KRT14 | hs00265033_m1 (TaqMan® probe ID) | |
| Differentiation Markers (qPCR) | NOGGIN | hs00271352_s1 (TaqMan® probe ID) | |
| Differentiation Markers (qPCR) | T | hs00610080_m1 (TaqMan® probe ID) | |
| Differentiation Markers (qPCR) | RUNX1 | hs01021970_m1 (TaqMan® probe ID) | |
| Differentiation Markers (qPCR) | DESMIN | hs00157258_m1 (TaqMan® probe ID) | |
| Differentiation Markers (qPCR) | PECAM1 | hs00169777_m1 (TaqMan® probe ID) | |
| Differentiation Markers (qPCR) | TAL1 | hs01097987_m1 (TaqMan® probe ID) | |
| Differentiation Markers (qPCR) | AFP | hs00173490_m1 (TaqMan® probe ID) | |
| Differentiation Markers (qPCR) | SOX17 | hs00751752_s1 (TaqMan® probe ID) | |
| Differentiation Markers (qPCR) | FOXA2 | hs00232764_m1 (TaqMan® probe ID) | |
| Differentiation Markers (qPCR) | SOX7 | hs00846731_s1 (TaqMan® probe ID) | |

**Supplementary Table 3**. Quality control measures applied to iPSCs.

| **Test Description** | **Method** | **Required Result** |
| --- | --- | --- |
| Cell Viability Assessment After Thawing | Colony doubling measurement | Colonies must form and double in diameter within 5 days |
| Sterility Testing | Culture testing using agar and broth media | Negative growth |
| Mycoplasma Detection | Quantitative RT-PCR analysis | Negative |
| Alkaline Phosphatase Detection | Cellular staining technique | Over 80% of cells must show positive staining |
| Sample Identity Verification | Short Tandem Repeat (STR) analysis | Must match the original parental cell profile |
| Viral and Transgene Analysis | Quantitative RT-PCR with specific primer sets | Must show no detection of SEV genome or transgenes |
| Surface Antigen (SA)  Expression of Stem Cell Markers | Combined immunostaining and flow cytometry | Over 80% of cells must express SSEA4 |
| Pluripotency Markers Analysis | Immunocytochemistry | Positive |
| Differentiation Capability | Embryoid body formation with gene expression analysis | Must show at least one gene per germ layer expressed at 2-fold or higher levels |
| Cytogenetic Analysis | G-banding technique | Normal karyotype (46 XX or 46 XY) |
